# Spatial epidemiology of gestational age and birth weight in Switzerland: Census-based linkage study

**DOI:** 10.1101/466854

**Authors:** Veronika Skrivankova, Marcel Zwahlen, Mark Adams, Nicola Low, Claudia E Kuehni, Matthias Egger

## Abstract

**Background:** Gestational age and birth weight are strong predictors of infant morbidity and mortality. Understanding spatial variation can inform policies to reduce health inequalities. We examined small-area variation in gestational age and birth weight in Switzerland.

**Methods:** All singleton live births recorded in the Swiss Live Birth Register 2011 to 2014 were eligible. We deterministically linked the Live Birth Register with census and survey data to create datasets including neonatal and pregnancy-related variables, parental characteristics and geographical variables. We produced maps of 705 areas and fitted linear mixed-effect models to assess to what extent spatial variation was explained by these variables.

**Results:** We analysed all 315,177 eligible live births. Area-level averages of gestational age varied between 272-279 days, and between 3138-3467g for birth weight. The fully adjusted models explained 31% and 87% of spatial variation of gestational age and birth weight, respectively. Language region explained most of the variation, with shorter gestational age and lower birth weight in French- and Italian- than in German-speaking areas. Other variables explaining variation were, for gestational age, the level of urbanisation, the parents’ nationality and missing father. For birth weight, they were gestational age, altitude, born out of wedlock, and parental nationality. In a subset of 69,463 live births with data on parental education, levels of education were only weakly associated with gestational age or birth weight.

**Conclusions:** In Switzerland, small area variation in birth weight is largely explained, and variation in gestational age partially explained by geocultural, socio-demographic and pregnancy factors.

**Strengths and limitations of this study:**

- This study was based on a large sample with national coverage, with data on neonatal and pregnancy-related predictors of gestational age and birth weight, and precise spatial data.
- No data were available on the mode of delivery, maternal smoking, mothers’ weight and height or gestational diabetes.
- The fully adjusted model explained about 80% of the regional variation in birth weight and about 40% of the variation in gestational age.
- Language region, a proxy for cultural, social and behavioural factors, was a strong explanatory factor, with lower birth weight and shorter gestation in the French and Italian compared to the German language region.
- Unknown father was associated with shorter gestation and lower birth weight, indicating that children not recognised by their fathers may be at higher risk of poor outcomes.

## INTRODUCTION

Gestational age and birth weight are important indicators of prenatal development and predictors of infant morbidity, mortality and long-term health [1–4]. An understanding of geographic differences and their determinants can help to develop policies that reduce health inequalities across population groups and regions [1–4]. Many genetic, physiological, pregnancy-related, socio-economic, lifestyle and environmental factors have been reported to influence gestational age and birth weight [5–8]. Some of these factors tend to cluster in space and regional differences in health outcomes may hence be partially explained by the spatial distribution of their predictors. Importantly, both individual-level factors and the social and environmental characteristics of communities and neighbourhoods may contribute to regional differences [9,10].

Variation across small areas in pregnancy outcomes have not been studied widely. In Scotland, small area crime rates were associated with lower birth weight and with the risk of both small for gestational age babies and preterm birth [11]. A study at county level in Georgia and South Carolina in the United States showed that the proportion of African Americans was associated with low birth weight, whereas higher income was associated with higher birth weight [12]. Similarly, neighbourhood racial composition contributed to variation in low birth weight in New York State [13]. Other small-area analyses have examined associations between birth outcomes and air pollution [14,15]. To our knowledge, few small-area analyses have considered gestational age.

In Switzerland, studies of pregnancy outcomes have focused on specific groups such as migrants or HIV-infected women [16,17], but have not examined geographic variations. The Federal Office of Statistics publishes routine statistics from the Live Birth Register, which does not include geographic information [18]. The objectives of this study were to conduct a nationwide analysis of spatial variation in gestational age and birth weight, and to assess how much small-area variation was explained by available data about neonatal and pregnancy-related variables, parental characteristics and geographical variables.

## METHODS

### Data sources

We used deterministic methods to link three data sources using encrypted national identification numbers: the Live Birth Register, the Swiss National Cohort and the Structural Surveys. Registration of live births is compulsory by law in Switzerland coverage is near 100%. The Swiss National Cohort (SNC) is a long-term, national study of mortality in Switzerland [19,20], linking census and mortality records. The 1990 and 2000 censuses were the last house-to-house censuses with coverage of the entire Swiss population. From 2010 onwards, the national census was replaced by a national population register and annual postal survey of the resident population, known as Structural Surveys [21]. Each structural Survey includes a random sample of around 300,000 people aged 15 years or older; for example, in 2010, it included 317,221 persons [21]. The reference is the entire Swiss resident population and the reference day 31 December.

### Variables and definitions

We defined three sets of variables. The first set, neonatal and pregnancy-related variables come from the Live Birth Register; date of birth, birth weight, gestational age, sex and birth rank. Birth weight is measured after initial mother-child bonding, usually by the midwife using a calibrated hospital scale. Gestational age is based on the last menstrual period, with or without additional information from ultrasound scans. Birth rank was classified as 1, 2, 3 and ≥4 live births, including the current birth. Birth rank is only available if the mother was married at the time of birth, and it is counted only within the current marriage. The second set includes parental variables. The Structural Surveys provide information about the highest level of completed maternal and paternal education, classified as ‘tertiary’, ‘secondary’, or ‘compulsory or less’. The Swiss National Cohort provides data about parental nationality categorised as ‘Swiss’, ‘Southern Europe’, ‘Western Europe’, ‘Northern Europe’, ‘Eastern Europe’, ‘Other’ (non-European), or missing (Supplementary Table S1 gives the full list of countries). The third set, geographical variables comes from the Swiss National Cohort. Each live birth was assigned an altitude and one of 705 statistical areas [22], based on the geocode of place of residence of the mother at the time of birth. Language regions are ‘German’, ‘French’ and ‘Italian’, and the level of urbanisation was defined using standard definitions of ‘urban’, ‘peri-urban’ and ‘rural’.

### Study populations and outcomes

All singleton live births recorded in the Live Birth Register from 1 January 2011 to 31 December 2014 were eligible. Gestational age at birth and birth weight were the outcomes of interest. For each outcome, two datasets were analysed: the first, larger dataset consisted of all eligible births with complete data on gestational age, birth weight and nationality of the mother. The second was the complete case population containing eligible live births with available data on all variables, including parental education. The second dataset included married mothers only who delivered at age 20 years or older because the birth rank is available for married women only, and education is incomplete below age 20 years.

### Statistical and spatial analyses

We fitted linear mixed-effect models (LMEM) to examine the associations between the two outcomes and the neonatal and pregnancy, parental and environmental factors. In the model for birth weight, we log-transformed the outcome and used a cubic spline function with three knots at weeks 25, 30 and 35 to capture the relationship between gestational age and log birth weight. Log transforming the birth weight results in a multiplicative model. Except for gestational age, maternal age and altitude, all predictors were modelled categorically. Maternal age was modelled by a piece-wise linear function, with age group 20 to 30 years as the reference group and separate linear trends for age groups 30-40 years, over 40 years and less than 20 years. Altitude was centred at 500 m and modelled linearly. The random effects in the mixed-effect model captured area-level differences between observed and expected mean outcome, based on the 705 statistical areas [22]. In the main analysis, we fitted four models to the complete-case dataset: Model 0 contained no explanatory variables. Model 1 included birth and pregnancy-related variables: sex, birth rank and gestational age (for the analysis of birth weight). Model 2 additionally included age of the mother, parental education and nationality. Model 3 additionally included geographical variables: altitude, degree of urbanisation and language region.

We displayed mean gestational age and birth weight at area-level on maps and assessed to what extent spatial variation was accounted for by the explanatory variables. Values were categorised into seven intervals symmetric around the mean and color-coded. Spatial autocorrelation of the gestational age and birth weight across regions was tested by global and local Moran’s I tests [23]. The global Moran test summarises overall spatial autocorrelation and the local test identifies areas that are correlated with neighbouring areas. In the presence of spatial autocorrelation, model estimates are at risk of bias if the autocorrelation is not taken into account.

In a sensitivity analysis, we accounted for spatial autocorrelation using the Besag-York-Mollier (BYM) model [24] using uninformative gamma-distributed (1, 0.005) priors. The calculations were carried out using the Integrated Nested Laplace Approximation (INLA) approach [25]. Similar results from models with and without the spatial component indicate low bias. Finally, we repeated analyses of birth weight without adjusting for gestational age. All analyses and maps were done in R 3.3.2 [26] using packages lme4, maptools, sp, spdep, rgdal, INLA, GISTools, rgeos, raster and ggplot2.

### Patient and public involvement

This analysis was based on routine registry data and no patients were involved in developing the research question, outcome measures and overall design of the study. Due to the anonymous nature of the data, we were unable to disseminate the results of the research directly to study participants.

## RESULTS

### Characteristics of study populations

A total of 328,349 live births were recorded in Switzerland between 1 January 2011 and 31 December 2014. We excluded non-singleton live births (n=11,835) and those with missing gestational age, birth weight or maternal nationality. The eligible study population therefore included 315,177 singleton live births. The complete case population consisted of 69,463 singleton live births with values available for all predictors including parental education, for which complete data were only available in the Structural Surveys (supplementary Figure S1).

Table 1 shows the distributions of predictors and outcomes in the two study populations. Data about the nationality of fathers was missing for 1.5% of eligible live births. In almost all of these cases, information about the father was missing completely, indicating that the father is unknown to the authorities. Apart from missing data, the distributions of most variables were similar between the two nested datasets. By design, the complete case population included married mothers only. The proportion of Swiss mothers and fathers was higher in the complete case population than in the eligible population. Birth at full term was defined as between 39 and 41 weeks of gestation (273 to 287 days). The mean gestational age in the eligible population was 276 days (SD 12) and the mean birth weight 3328 g (SD 515). The corresponding figures in the complete case population were 276 days (SD 12) and 3349 g (SD 501).

**Table 1.** Characteristics of complete case and eligible study populations.

|  | Eligible population |  |  | Complete case population |  |  |
| --- | --- | --- | --- | --- | --- | --- |
|  | No. (%) | Gestational age (days)<br>Mean (SD) | Birth weight (g)<br>Mean (SD) | No. (%) | Gestational age (days)<br>Mean (SD) | Birth weight (g)<br>Mean (SD) |
| **Total** | 315177 (100%) | 276 (12) | 3328 (515) | 69463 (100%) | 276 (12) | 3349 (501) |
| **Birth weight (g)** |  |  |  |  |  |  |
| <1500 | 2141 (0.7%) | 196 (27) | 966 (354) | 347 (0.5%) | 197 (29) | 969 (516) |
| 1500-1999 | 2413 (0.8%) | 238 (15) | 1800 (142) | 513 (0.7%) | 239 (15) | 1804 (503) |
| 2000-2499 | 10036 (3.2%) | 258 (14) | 2312 (134) | 1993 (2.9%) | 258 (13) | 2313 (490) |
| ≥ 2500 | 300586 (95.4%) | 277 (9) | 3391 (423) | 66609 (95.9%) | 277 (9) | 3404 (502) |
| **Gestation. age (weeks)** |  |  |  |  |  |  |
| <32<sup>0</sup> | 2333 (0.7%) | 195 (23) | 1108 (527) | 381 (0.5%) | 195 (24) | 1127 (512) |
| 32<sup>0</sup>-34<sup>6</sup> | 3950 (1.3%) | 237 (6) | 2144 (424) | 800 (1.2%) | 237 (6) | 2138 (492) |
| 35<sup>0</sup>-36<sup>6</sup> | 10907 (3.5%) | 253 (4) | 2686 (431) | 2296 (3.3%) | 253 (4) | 2698 (490) |
| ≥ 37<sup>0</sup> | 297987 (94.5%) | 278 (8) | 3385 (440) | 65986 (95%) | 278 (8) | 3399 (502) |
| **Sex** |  |  |  |  |  |  |
| Female | 152757 (48.5%) | 276 (12) | 3260 (494) | 33698 (48.5%) | 276 (11) | 3277 (499) |
| Male | 162420 (51.5%) | 275 (13) | 3392 (525) | 35765 (51.5%) | 276 (12) | 3416 (504) |
| **Birth rank** |  |  |  |  |  |  |
| 1 | 115871 (36.8%) | 276 (13) | 3278 (511) | 30647 (44.1%) | 276 (12) | 3278 (503) |
| 2 | 97705 (31%) | 275 (11) | 3390 (493) | 28544 (41.1%) | 276 (11) | 3389 (502) |
| 3 | 28738 (9.1%) | 275 (11) | 3433 (502) | 8154 (11.7%) | 276 (11) | 3438 (501) |
| ≥ 4 | 7616 (2.4%) | 276 (12) | 3463 (527) | 2118 (3%) | 276 (11) | 3479 (494) |
| missing (not married) | 65247 (20.7%) | 276 (14) | 3263 (536) | - | - | - |
| **Civil status** |  |  |  |  |  |  |
| Married | 250055 (79.3%) | 276 (12) | 3345 (508) | 69463 (100%) | 276 (12) | 3349 (502) |
| Single | 56462 (17.9%) | 276 (14) | 3263 (533) | 0 (0%) | - | - |
| Divorced | 8353 (2.7%) | 274 (14) | 3258 (556) | 0 (0%) | - | - |
| Widow | 307 (0.1%) | 274 (14) | 3286 (550) | 0 (0%) | - | - |
| **Maternal age (years)** |  |  |  |  |  |  |
| mean (SD) | 31.7 (5.0) |  |  | 32.2 (4.6) |  |  |
| < 20 | 2679 (0.8%) | 275 (16) | 3224 (554) | 0 (0%) | - | - |
| ≥ 20-25 | 28615 (9.1%) | 277 (12) | 3317 (511) | 4365 (6.3%) | 277 (12) | 3359 (497) |
| ≥ 25-30 | 82620 (26.2%) | 276 (12) | 3330 (506) | 17764 (25.6%) | 276 (11) | 3346 (505) |
| ≥ 30-35 | 118303 (37.5%) | 276 (12) | 3335 (510) | 28108 (40.5%) | 276 (12) | 3351 (501) |
| ≥ 35-40 | 67914 (21.5%) | 275 (12) | 3333 (523) | 16021 (23.1%) | 275 (11) | 3353 (501) |
| ≥ 40 | 15046 (4.8%) | 273 (14) | 3286 (555) | 3205 (4.6%) | 273 (14) | 3304 (501) |
| **Nationality mother** |  |  |  |  |  |  |
| Switzerland | 194570 (61.7%) | 276 (12) | 3322 (511) | 46651 (67.2%) | 276 (11) | 3342 (504) |
| Southern Europe | 23585 (7.5%) | 275 (12) | 3251 (494) | 4763 (6.9%) | 276 (11) | 3269 (501) |
| Western Europe | 26005 (8.3%) | 276 (12) | 3348 (516) | 4799 (6.9%) | 276 (12) | 3369 (499) |
| Northern Europe | 3695 (1.2%) | 276 (13) | 3418 (510) | 703 (1%) | 276 (13) | 3433 (489) |
| Eastern Europe | 38762 (12.3%) | 276 (13) | 3397 (523) | 7743 (11.1%) | 276 (12) | 3428 (499) |
| Other | 28560 (9.1%) | 275 (14) | 3313 (535) | 4804 (6.9%) | 275 (13) | 3331 (492) |
| **Nationality father** |  |  |  |  |  |  |
| Switzerland | 191589 (60.8%) | 276 (12) | 3329 (506) | 47018 (67.7%) | 276 (12) | 3346 (504) |
| Southern Europe | 31466 (10%) | 275 (12) | 3256 (493) | 6473 (9.3%) | 275 (11) | 3273 (499) |
| Western Europe | 26954 (8.6%) | 276 (12) | 3353 (518) | 4949 (7.1%) | 276 (12) | 3376 (486) |
| Northern Europe | 3911 (1.2%) | 276 (12) | 3406 (510) | 724 (1%) | 276 (13) | 3412 (492) |
| Eastern Europe | 35387 (11.2%) | 276 (13) | 3397 (528) | 6960 (10%) | 276 (12) | 3424 (503) |
| Other | 21077 (6.7%) | 276 (13) | 3307 (531) | 3339 (4.8%) | 276 (12) | 3318 (502) |
| missing | 4793 (1.5%) | 272 (23) | 3148 (693) | - | - | - |
| **Education mother** |  |  |  |  |  |  |
| Tertiary | 42088 (13.4%) | 276 (12) | 3344 (500) | 28016 (40.3%) | 276 (12) | 3356 (499) |
| Secondary | 48878 (15.5%) | 276 (12) | 3328 (509) | 32614 (47.0%) | 276 (12) | 3343 (505) |
| Compulsory | 14642 (4.6%) | 275 (13) | 3329 (534) | 8833 (12.7%) | 275 (12) | 3345 (501) |
| Unknown (age <20 yrs) | 2679 (0.8%) | 275 (16) | 3224 (554) | - | - | - |
| missing | 206890 (65.6%) | 276 (12) | 3326 (517) | - | - | - |
| **Education father** |  |  |  |  |  |  |
| Tertiary | 49848 (15.8%) | 276 (12) | 3348 (497) | 34325 (49.4%) | 276 (12) | 3357 (501) |
| Secondary | 41301 (13.1%) | 276 (12) | 3323 (511) | 26857 (38.7%) | 276 (12) | 3340 (502) |
| Compulsory | 13731 (4.4%) | 276 (12) | 3323 (514) | 8281 (11.9%) | 275 (12) | 3340 (506) |
| missing | 210297 (66.7%) | 276 (13) | 3325 (519) | - | - | - |
| **Altitude (m)** |  |  |  |  |  |  |
| mean (SD) | 515 (189) |  |  | 511 (181) |  |  |
| **Urbanisation** |  |  |  |  |  |  |
| Urban | 96643 (30.7%) | 276 (13) | 3326 (517) | 18516 (26.7%) | 276 (12) | 3344 (498) |
| Peri-urban | 138826 (44%) | 275 (12) | 3329 (514) | 31430 (45.2%) | 276 (12) | 3348 (501) |
| Rural | 79708 (25.3%) | 276 (12) | 3329 (512) | 19517 (28.1%) | 276 (12) | 3354 (506) |
| **Language region** |  |  |  |  |  |  |
| German | 223586 (70.9%) | 276 (12) | 3348 (515) | 46546 (67%) | 276 (12) | 3370 (502) |
| French | 80068 (25.4%) | 275 (12) | 3283 (512) | 19324 (27.8%) | 275 (11) | 3310 (502) |
| Italian | 11523 (3.7%) | 275 (12) | 3252 (494) | 3593 (5.2%) | 275 (12) | 3273 (494) |

### Maps of gestational age and birth weight

Figure 1 presents maps of Switzerland with crude average gestational age and birth weight across the 705 areas. For both outcomes, the maps are broadly similar between the eligible and complete case populations. For gestational age, area-level averages for the eligible population vary between 272 and 279 days. For the complete case population variation was greater, from 265 to 281 days, as expected for a smaller sample. The map shows shorter gestation in the Western, North Western region and Southern (Canton of Ticino) regions of Switzerland, with a patchy pattern in the densely populated areas between the Alps (across the centre) and Jura mountain ranges (to the North West). For birth weight, area-level averages vary between 3138 and 3467g for the eligible population and between 3020 and 3597g for the complete case population. The maps for birth weight show lower birth weights in the Western and Southern regions of the country. The French and Italian-speaking regions are in the West and South of Switzerland, with the remainder being German-speaking.

**Figure 1.**
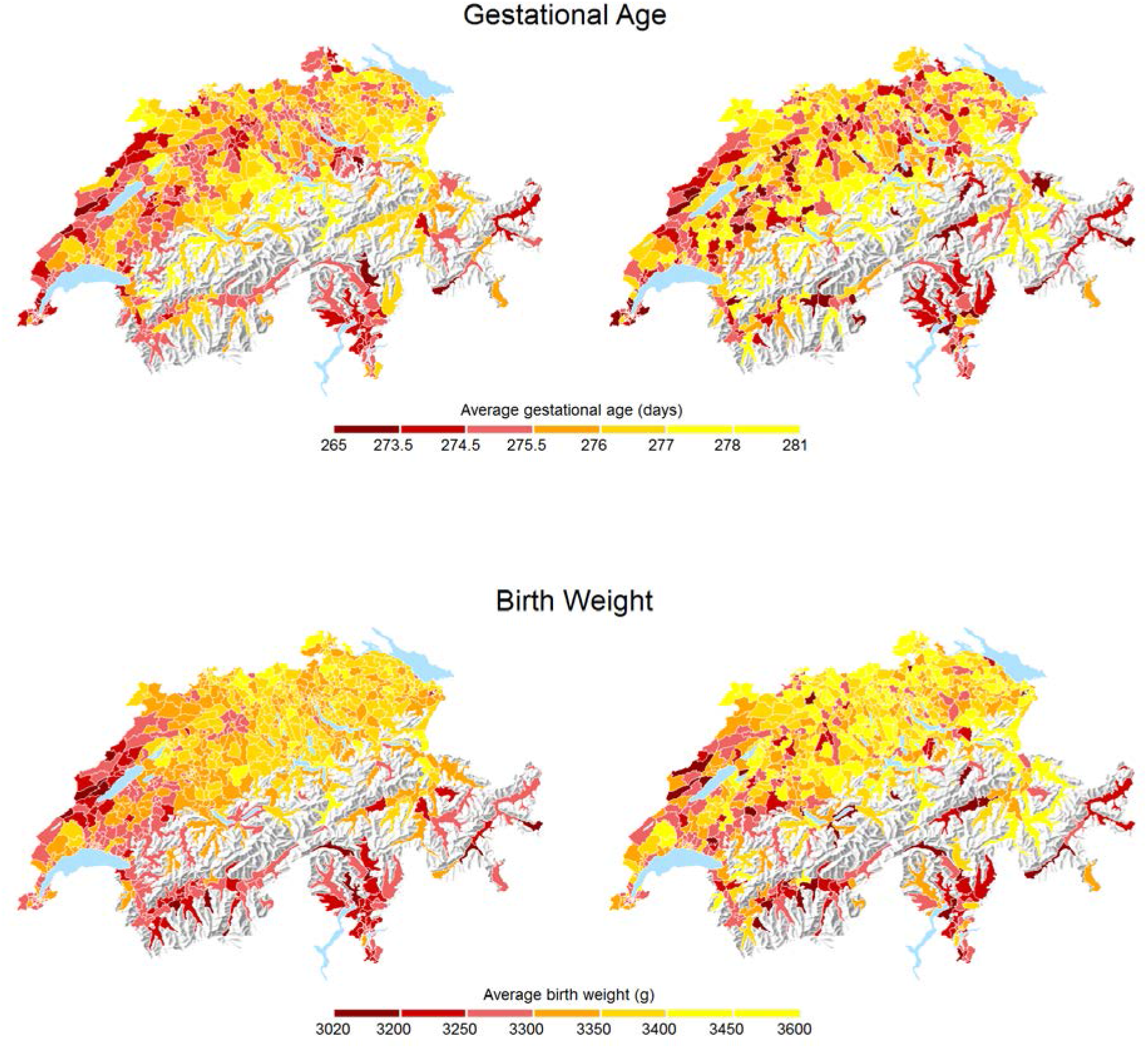
Maps of average gestational age (upper two panels) and birth weight (lower two panels) observed across 705 Swiss areas. Left: all eligible live births (n=315,177), right: complete case population (n=69,463).

### Multivariable analyses

Table 2 shows associations of area-level mean gestational age at birth and mean birth weight with pregnancy, parental and environmental factors from the fully adjusted linear mixed-effects models (model 3). For gestational age, the largest differences are observed across categories of maternal age at birth, with pregnancies in mothers aged 40 years or older, and below 20 years about 3 days shorter than in mothers aged 20 to 30 years in both the eligible and the complete case populations. Of note, compared with Swiss fathers, pregnancies were about 4 days shorter if the nationality of the father was missing. Smaller differences in gestational age were observed across categories of sex, birth rank, nationality of the mother, urbanisation and between language regions (Table 2). In the complete case population, lower levels of education were associated with shorter pregnancies. Gestational age at birth was not associated with altitude.

**Table 2.**
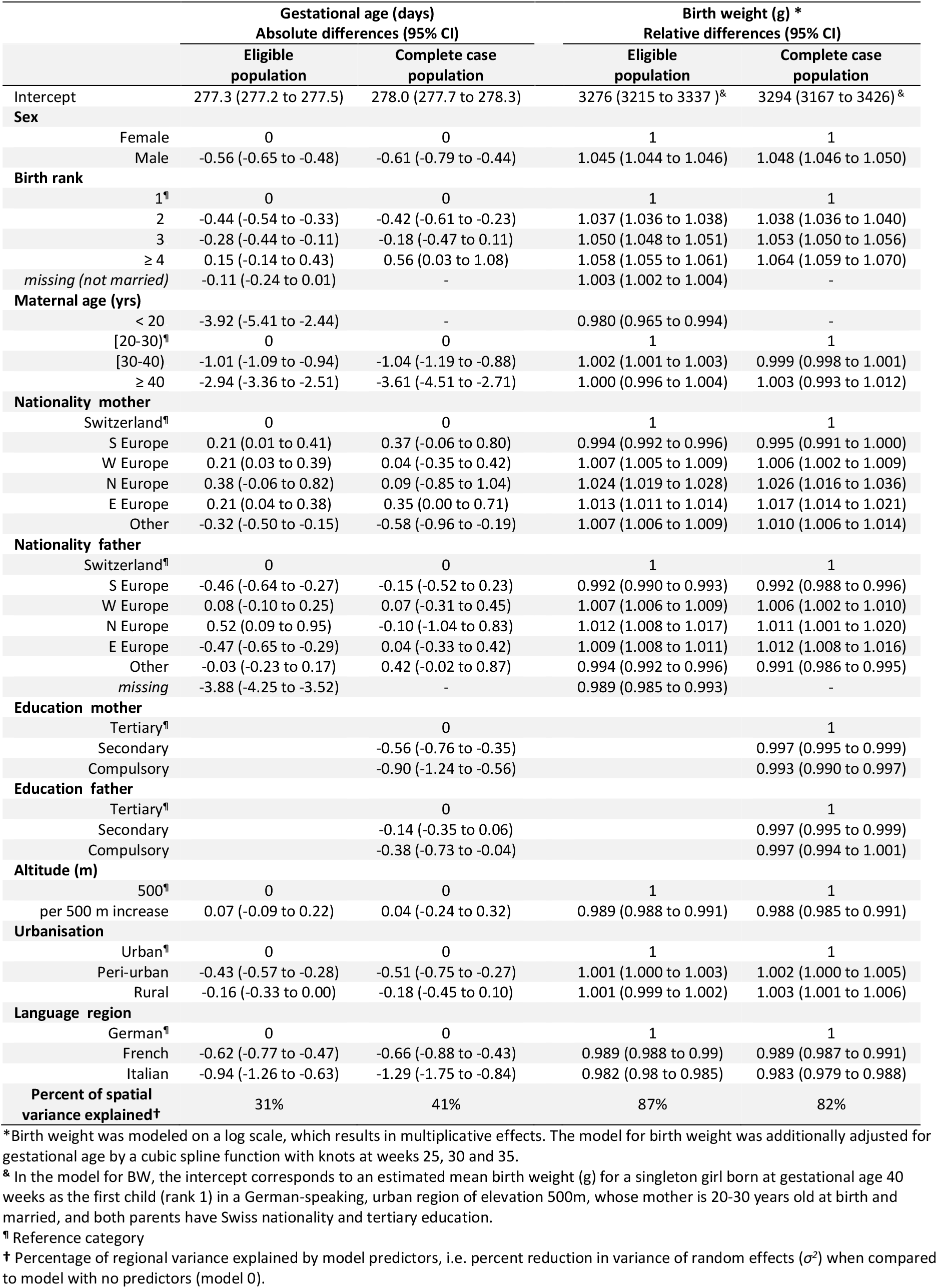
Associations of mean gestational age at birth and mean birth weight with pregnancy, parental and environmental factors from adjusted linear mixed-effects model (model 3).

Supplementary Figure S2 shows the relationship between gestational age and birth weight separately for male and female newborns. Male newborns were about 5% heavier than female newborns and birth weight increased with birth order (Table 2). In contrast to gestational age, mother’s age was not associated with birth weight. Babies born to mothers or fathers from Northern or Eastern Europe were slightly heavier than babies born to Swiss mothers; birth weights were lowest for babies of fathers with missing nationality. Birth weight slightly decreased with increasing parental educational attainment. Babies born in the French and Italian-speaking regions were lighter than babies born in the German-speaking Switzerland. Finally, birth weight decreased with increasing altitude of residence.

### Proportion of spatial variation explained

The fully adjusted model (model 3) for gestational age explained 31% and 41% of the spatial variation across the 705 areas for eligible and complete case populations, respectively. The corresponding figures for birth weight were 87% and 82% (Figure 2). When assessing each factor separately (Table 3), language region alone explained most of the spatial variation for both outcomes. For gestational age, level of urbanisation of the mother’s place of residence also explained part of the variation. Factors that also contributed to explaining the spatial variation in birth weight were gestational age, parental nationalities, altitude at the mother’s place of residence and birth order. Figure 3 illustrates the reduction in the spatial variation of gestational age and birth weight with maps, when moving from model 0 (0% reduction) to models 1, 2 and 3, based on the complete case population.

**Table 3.**
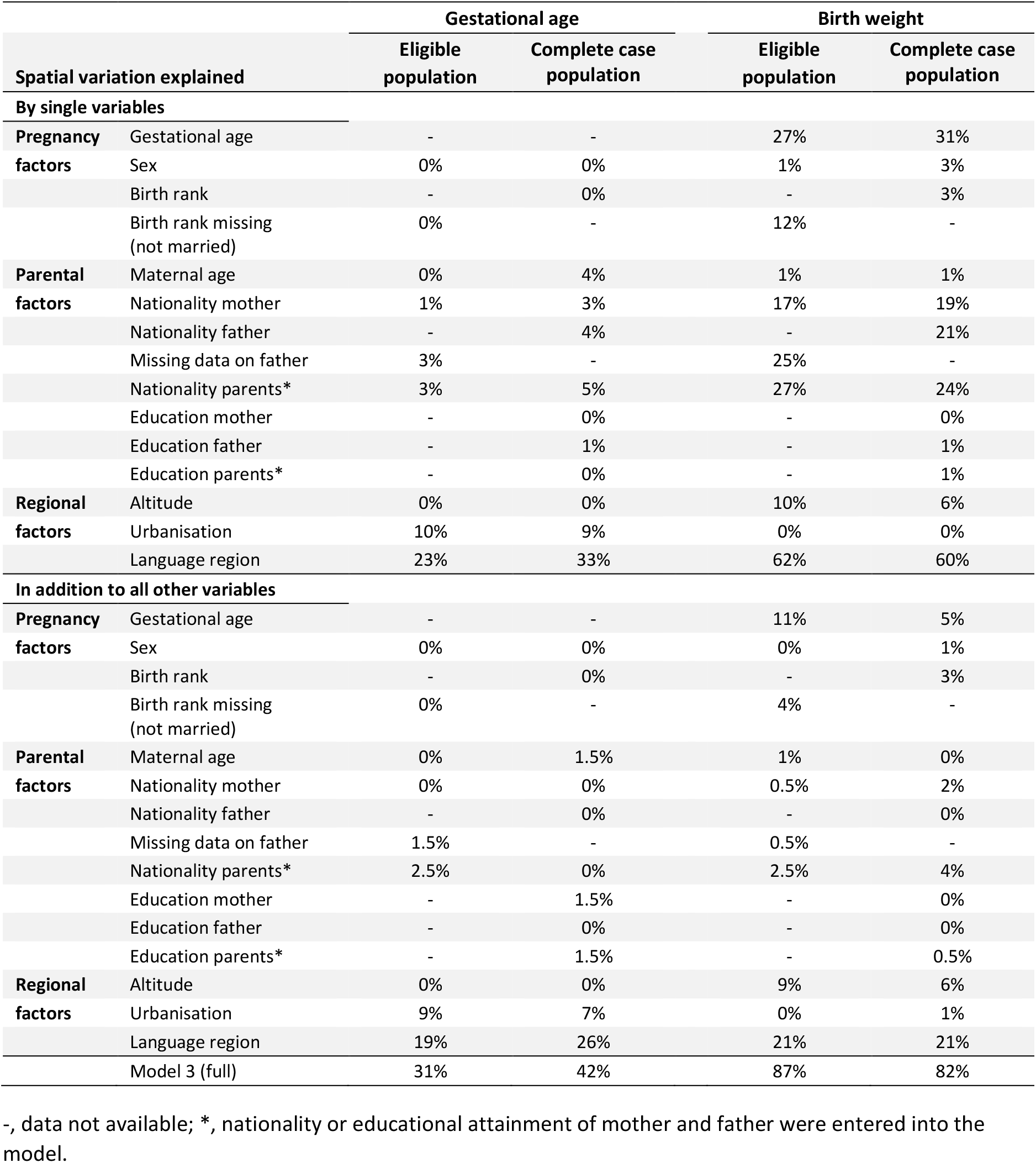
Percentage of spatial variation explained by each individual variable and explained in addition after adjusting for all other variables.

**Figure 2.**
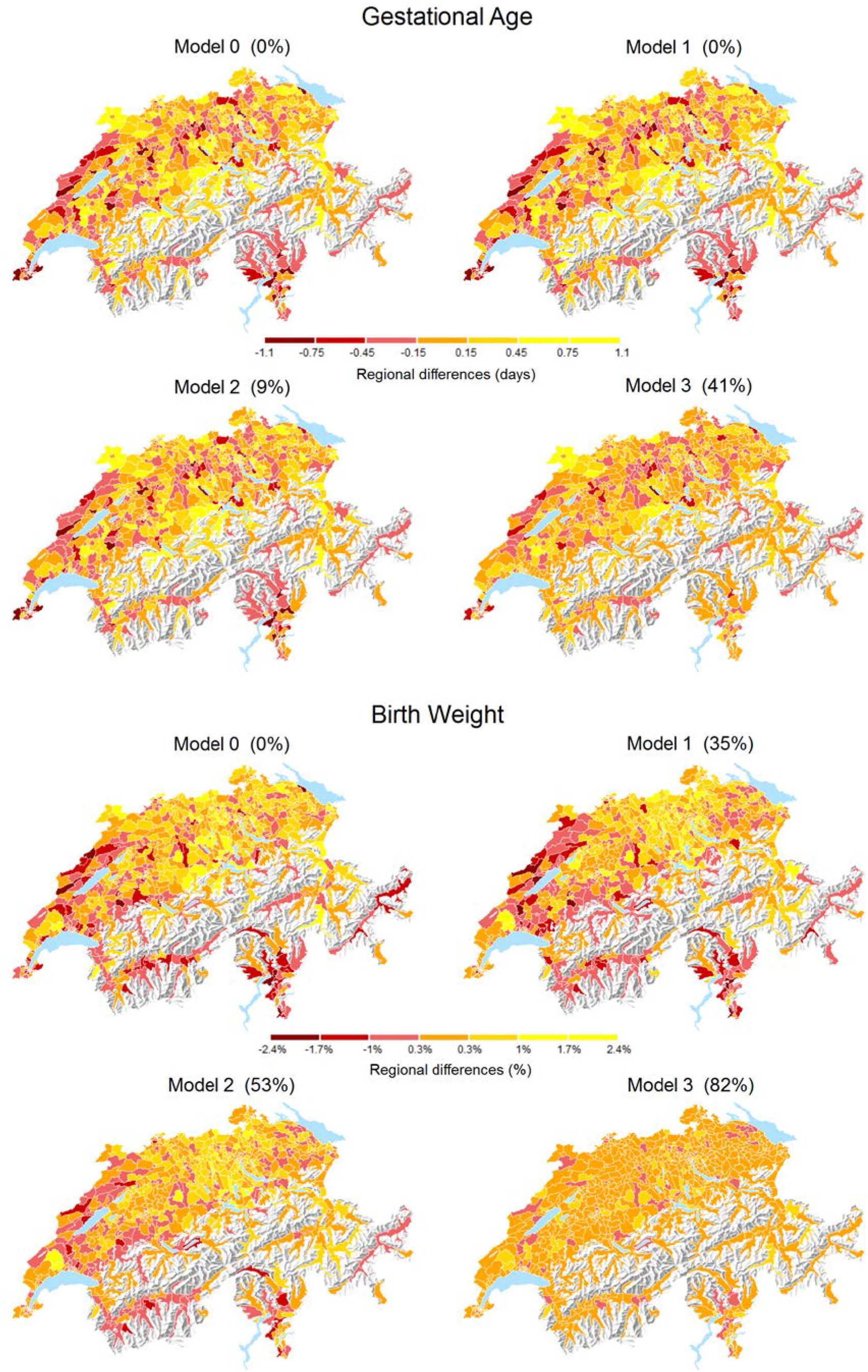
Maps of gestational age and birth weight from crude model (model 0) and multivariable linear mixed-effect models (models 1-3) with percent reduction in the regional variation, represented by random effects. Analyses based on complete case population (*N* = 69,463).

### Spatial autocorrelation and sensitivity analyses

For gestational age, the global Moran’s I statistic, based on the complete case dataset and model 0, was I=0.19, with P<10^-13^. After adjusting for all the predictors in model 3 there was still some residual autocorrelation (*I*=0.09, P=0.0001). For birth weight, the corresponding Moran’s I statistic was I=0.26, with P<10^-15^. After adjusting for all predictors in model 3 there was little residual autocorrelation (I=0.04, P=0.07). Supplementary Table S2 compares the results from model 3 accounting and not accounting for spatial autocorrelation. The results are similar and the potential bias from residual spatial autocorrelation is therefore unlikely to be a major issue. Repeating analyses of birth weight without adjusting for gestational age produced generally similar coefficients (Table S3). Associations with maternal age, maternal education and language regions were slightly stronger in model 3 without adjustment for gestational age, possibly because some of their effect was mediated by gestational age. Model 3 without gestational age explained 77% of the spatial variation both in the eligible and complete case populations.

## DISCUSSION

Our study assessed factors associated with gestational age and birth weight in Switzerland and their contribution to spatial variation, based on routinely collected data. Gestational age at birth was strongly associated with maternal age, missing information on the father and language region. Birth weight was associated with sex, birth rank, missing information on the father, parental education, altitude and language region. There was substantial regional variation and spatial autocorrelation across regions. The variables included in the fully adjusted model explained about 80% of the regional variation in birth weight and about 40% of the regional variation in gestational age. Strengths of this study include a large sample with national coverage of the Swiss resident population, as well as the availability of data on several relevant predictors, either on all births or on a large random sample of eligible births. Precise spatial data and spatial statistics allowed us to assess the proportion of area-level variation explained, spatial autocorrelation and gauge the likelihood of bias due to residual autocorrelation.

This study found important spatial variation in both gestational age and birth weight in Switzerland. Language region in Switzerland was the single factor that explained the greatest proportion of spatial variation in gestational age and birth weight. In the French and Italian speaking regions, gestational age was shorter and birth weight lower than in the German speaking part. Language region combines a wide range of cultural, social and behavioural factors, including diet, smoking and alcohol consumption [27] of parents, as well as their ancestry, which probably explain its strong explanatory power. Other factors that could not be measured directly, such as health care provision, might have accounted for some of the unexplained variation. Data about the mode of delivery (vaginal or by Caesarean section, induced or spontaneous) were not available. Whilst Caesarean section rates vary geographically, they are unlikely to account for the observed spatial variation in gestational age at birth. Geographical patterns of Caesarean section are largely driven by urban-rural differences [28].

While young and old maternal age are well-known predictors of shorter gestation [29,30], the association we found with missing data on the father’s nationality was somewhat unexpected. In the vast majority of cases, the information is missing because no father came forward and officially accepted paternity of the child. It is possible that missing data about the father are an indicator of lower socio-economic position and social support of the mother, resulting in greater vulnerability. Studies from the United States of America found a missing name of the father on the infant’s birth certificate was associated with lower education, smoking during pregnancy, preterm birth, lower birth weight, no breastfeeding and higher neonatal and post-neonatal mortality [31–34]. Children not recognised by their fathers may thus be a group at higher risk of infant and child morbidity and mothers might benefit from additional care during pregnancy and postnatally.

There are several limitations to our study. The complete case dataset was restricted to married mothers because the Swiss Live Birth Register only records birth rank if the mother was married at the time of birth. This limitation might have resulted, for example, in the weaker than expected association between birth weight and parental education. Studies from countries such as the Netherlands have shown larger gaps across levels of educational attainment, which were largest amongst unmarried women [35]. We did not have data about maternal health-related behaviours such as smoking [36], mothers’ weight and height [36], disease such as gestational diabetes and data on parental genetic factors. Whilst parental nationality and education might have served as crude proxies for some missing variables, individual-level data about these factors would be valuable. A recent large-scale meta-analysis of genome-wide association data indicated that genetic factors influence birth weight through their effects on gestational age, maternal glucose metabolism, cytochrome P450 activity and possibly on maternal immune function and blood pressure [37]. Of note, compared to the foetus who carries maternal and paternal genes, maternal genes exert a larger effect on gestational age and a weaker effect on birth weight [38,39]. Examining the proportion of preterm births (before 37 weeks) or the proportion of low birth weights (<2500g) might seem clinically more relevant than the means examined in this study. However, from a statistical point of view, dichotomizing continuous data is “a practice to avoid” [40], while the mean observed in a region and the proportion of preterm and low birth weight births are highly correlated, as shown in supplementary Figure S3.

We adjusted analyses of birth weight for gestational age, which may mediate the effects of other variables, for example maternal age. Adjusting for a variable on the causal pathway has been criticised because it may introduce selection bias (or collider bias in the language of directed acyclic graphs), if there are unknown or unmeasured factors that have an effect on both gestational age and birth weight [41–43]. In our study results were broadly similar with and without adjustment for gestational age and the focus of our study was not on causal inference, but on gaining an understanding of the factors contributing to spatial variation of birth weight and gestational age.

In conclusion, our study identified important differences in mean gestational age and birth weight across Switzerland. Small area variation in birth weight is largely, and in gestational age partially, explained by pregnancy-related, parental, and environmental factors.

## SUPPLEMENTARY MATERIALS

**Supplementary Table S1.** Number of live births, mean gestational age and mean birth weight by maternal nationality in the eligible population (*N* = 315’177).

| Nationality | N | Gestational age (days)<br>Mean (SD) | Birth weight (g)<br>Mean (SD) |
| --- | --- | --- | --- |
| Switzerland | 194,570 | 276 (12) | 3322 (511) |
| **Southern Europe** |  |  |  |
| Andorra | 1 | 279 (-) | 3080 (-) |
| Italy | 8337 | 275 (12) | 3271 (496) |
| Malta | 13 | 273 (8) | 3188 (427) |
| Portugal | 12,368 | 276 (12) | 3235 (493) |
| San Marino | 2 | 274 (1.4) | 3485 (120) |
| Spain | 2864 | 276 (12) | 3263 (488) |
| **Western Europe** |  |  |  |
| Austria | 1555 | 275 (14) | 3328 (528) |
| Belgium | 583 | 276 (12) | 3357 (482) |
| Germany | 16,736 | 276 (13) | 3369 (517) |
| France | 6173 | 276 (12) | 3294 (505) |
| Lichtenstein | 100 | 275 (11) | 3369 (488) |
| Luxembourg | 55 | 276 (19) | 3396 (636) |
| Netherlands | 803 | 276 (12) | 3377 (529) |
| **Northern Europe** |  |  |  |
| Denmark | 271 | 276 (13) | 3383 (511) |
| Estonia | 81 | 279 (7) | 3601 (466) |
| Finland | 312 | 276 (11) | 3465 (523) |
| Ireland | 212 | 276 (16) | 3446 (548) |
| Iceland | 31 | 272 (25) | 3180 (775) |
| Latvia | 187 | 279 (9) | 3493 (434) |
| Lithuania | 152 | 277 (13) | 3450 (535) |
| Norway | 110 | 275 (11) | 3390 (525) |
| Sweden | 571 | 276 (12) | 3422 (470) |
| UK | 1768 | 276 (13) | 3397 (513) |
| **Eastern Europe** |  |  |  |
| Czech Republic | 623 | 275 (12) | 3339 (499) |
| Hungary | 913 | 275 (13) | 3341 (512) |
| Poland | 1778 | 276 (12) | 3399 (497) |
| Slovakia | 1068 | 276 (12) | 3348 (509) |
| Albania | 209 | 276 (12) | 3406 (476) |
| Bosnia & Herzegovina | 1952 | 276 (12) | 3466 (492) |
| Croatia | 1582 | 276 (12) | 3448 (540) |
| Kosovo | 10,278 | 276 (13) | 3421 (530) |
| Macedonia | 5842 | 276 (13) | 3392 (514) |
| Montenegro | 212 | 276 (10) | 3416 (466) |
| Serbia | 5195 | 276 (13) | 3400 (536) |
| Serbia & Montenegro | 10 | 277 (8) | 3637 (250) |
| Slovenia | 163 | 275 (15) | 3366 (589) |
| Cyprus | 15 | 278 (8) | 3411 (525) |
| Bulgaria | 406 | 273 (15) | 3291 (559) |
| Greece | 375 | 274 (13) | 3317 (516) |
| Romania | 971 | 274 (14) | 3284 (537) |
| Turkey | 4441 | 275 (13) | 3347 (523) |
| Belarus | 172 | 277 (13) | 3385 (508) |
| Moldova | 135 | 276 (10) | 3496 (515) |
| Russia | 1567 | 277 (12) | 3427 (513) |
| Ukraine | 855 | 277 (11) | 3412 (473) |
| **Other (non-Europe)** |  |  |  |
| 6 most numerous: |  |  |  |
| Eritrea | 2600 | 279 (14) | 3380 (528) |
| Brazil | 2381 | 274 (12) | 3312 (498) |
| Sri Lanka | 1391 | 273 (14) | 3158 (553) |
| USA | 1291 | 276 (14) | 3378 (532) |
| China | 1293 | 276 (13) | 3425 (541) |
| Morocco | 1159 | 276 (14) | 3378 (536) |
| ... |  |  |  |
| **Total** | **315,177** | **276 (12)** | **3328 (515)** |

**Supplementary Table S2.** Comparison of results from fully adjusted model (model 3) accounting and not accounting for spatial autocorrelation. Based on complete-case population (*N* = 69′463).

|  | Accounting for spatial autocorrelation |  | Not accounting for spatial autocorrelation |  |
| --- | --- | --- | --- | --- |
|  | Gestational age (days)<br>Absolute differences<br>(95% CI) | Birth weight (g) *<br>Relative differences<br>(95% CI) | Gestational age (days)<br>Absolute differences<br>(95% CI) | Birth weight (g) *<br>Relative differences<br>(95% CI) |
| Intercept | 277.9 (277.6, 278.2) | 3289 (3159, 3423) | 278.0 (277.7 to 278.3) | 3294 (3167 to 3426) <sup>&</sup> |
| **Sex** |  |  |  |  |
| Female | 0 | 1 | 0 | 1 |
| Male | -0.61 (-0.79, -0.44) | 1.048 (1.046, 1.049) | -0.61 (-0.79 to -0.44) | 1.048 (1.046 to 1.050) |
| **Birth rank** |  |  |  |  |
| 1<sup>†</sup> | 0 | 1 | 0 | 1 |
| 2 | -0.42 (-0.61, -0.23) | 1.038 (1.036, 1.040) | -0.42 (-0.61 to -0.23) | 1.038 (1.036 to 1.040) |
| 3 | -0.18 (-0.47, 0.11) | 1.053 (1.050, 1.056) | -0.18 (-0.47 to 0.11) | 1.053 (1.050 to 1.056) |
| ≥ 4 | 0.55 (0.03, 1.08) | 1.064 (1.059, 1.070) | 0.56 (0.03 to 1.08) | 1.064 (1.059 to 1.070) |
| **Maternal age (yrs)** |  |  |  |  |
| [20-30)<sup>‡</sup> | 0 | 1 | 0 | 1 |
| [30-40) | -1.03 (-1.19, -0.87) | 0.999 (0.997, 1.001) | -1.04 (-1.19 to -0.88) | 0.999 (0.998 to 1.001) |
| ≥ 40 | -3.61 (-4.52, -2.71) | 1.003 (0.993, 1.012) | -3.61 (-4.51 to -2.71) | 1.003 (0.993 to 1.012) |
| **Nationality mother** |  |  |  |  |
| Switzerland<sup>‡</sup> | 0 | 1 | 0 | 1 |
| S Europe | 0.37 (-0.07, 0.80) | 0.995 (0.991, 1.000) | 0.37 (-0.06 to 0.80) | 0.995 (0.991 to 1.000) |
| W Europe | 0.03 (-0.35, 0.42) | 1.005 (1.001, 1.009) | 0.04 (-0.35 to 0.42) | 1.006 (1.002 to 1.009) |
| N Europe | 0.10 (-0.85, 1.05) | 1.025 (1.015, 1.035) | 0.09 (-0.85 to 1.04) | 1.026 (1.016 to 1.036) |
| E Europe | 0.35 (0.00, 0.71) | 1.017 (1.013, 1.021) | 0.35 (0.00 to 0.71) | 1.017 (1.014 to 1.021) |
| Other | -0.56 (-0.95, -0.18) | 1.010 (1.006, 1.014) | -0.58 (-0.96 to -0.19) | 1.010 (1.006 to 1.014) |
| **Nationality father** |  |  |  |  |
| Switzerland<sup>‡</sup> | 0 | 1 | 0 | 1 |
| S Europe | -0.14 (-0.52, 0.23) | 0.992 (0.988, 0.995) | -0.15 (-0.52 to 0.23) | 0.992 (0.988 to 0.996) |
| W Europe | 0.07 (-0.31, 0.45) | 1.006 (1.002, 1.010) | 0.07 (-0.31 to 0.45) | 1.006 (1.002 to 1.010) |
| N Europe | -0.08 (-1.02, 0.85) | 1.010 (1.001, 1.020) | -0.10 (-1.04 to 0.83) | 1.011 (1.001 to 1.020) |
| E Europe | 0.04 (-0.34, 0.42) | 1.011 (1.007, 1.015) | 0.04 (-0.33 to 0.42) | 1.012 (1.008 to 1.016) |
| Other | 0.44 (-0.01, 0.89) | 0.991 (0.986, 0.995) | 0.42 (-0.02 to 0.87) | 0.991 (0.986 to 0.995) |
| **Education mother** |  |  |  |  |
| Tertiary<sup>‡</sup> | 0 | 1 | 0 | 1 |
| Secondary | -0.56 (-0.77, -0.36) | 0.997 (0.995, 0.999) | -0.56 (-0.76 to -0.35) | 0.997 (0.995 to 0.999) |
| Compulsory | -0.91 (-1.25, -0.57) | 0.994 (0.990, 0.997) | -0.90 (-1.24 to -0.56) | 0.993 (0.990 to 0.997) |
| **Education father** |  |  |  | * |
| Tertiary<sup>‡</sup> | 0 | 1 | 0 | 1 |
| Secondary | -0.14 (-0.35, 0.06) | 0.997 (0.995, 0.999) | -0.14 (-0.35 to 0.06) | 0.997 (0.995 to 0.999) |
| Compulsory | -0.39 (-0.73, -0.04) | 0.997 (0.994, 1.001) | -0.38 (-0.73 to -0.04) | 0.997 (0.994 to 1.001) |
| **Altitude (m)** |  |  |  |  |
| 500<sup>‡</sup> | 0 | 1 | 0 | 1 |
| per 500 m increase | -0.01 (-0.31, 0.29) | 0.989 (0.985, 0.993) | 0.04 (-0.24 to 0.32) | 0.988 (0.985 to 0.991) |
| **Urbanisation** |  |  |  |  |
| Urban<sup>‡</sup> | 0 | 1 | 0 | 1 |
| Peri-urban | -0.49 (-0.73, -0.25) | 1.001 (0.998, 1.004) | -0.51 (-0.75 to -0.27) | 1.002 (1.000 to 1.005) |
| Rural | -0.19 (-0.47, 0.09) | 1.002 (0.998, 1.006) | -0.18 (-0.45 to 0.10) | 1.003 (1.001 to 1.006) |
| **Language region** |  |  |  |  |
| German<sup>‡</sup> | 0 | 1 | 0 | 1 |
| French | -0.41 (-0.77, -0.05) | 0.992 (0.984, 1.000) | -0.66 (-0.88 to -0.43) | 0.989 (0.987 to 0.991) |
| Italian | -1.29 (-1.75, -0.83) | 0.984 (0.978, 0.990) | -1.29 (-1.75 to -0.84) | 0.983 (0.979 to 0.988) |

\*Birth weight was modeled on a log scale, which results in multiplicative effects. The model for birth weight was additionally adjusted for gestational age by a cubic spline function with knots at weeks 25, 30 and 35.
<sup>&</sup> In the model for BW, the intercept corresponds to an estimated mean birth weight (g) for a singleton girl born at gestational age 40 weeks as the first child (rank 1) in a German-speaking, urban region of elevation 500m, whose mother is 20-30 years old at birth and married, and both parents have Swiss nationality and tertiary education.
<sup>‡</sup> Reference category
<sup>†</sup> Percentage of regional variance explained by model predictors, i.e. percent reduction in variance of random effects ( $\sigma^2$ ) when compared to model with no predictors (model 0).

**Supplementary Table S3.**
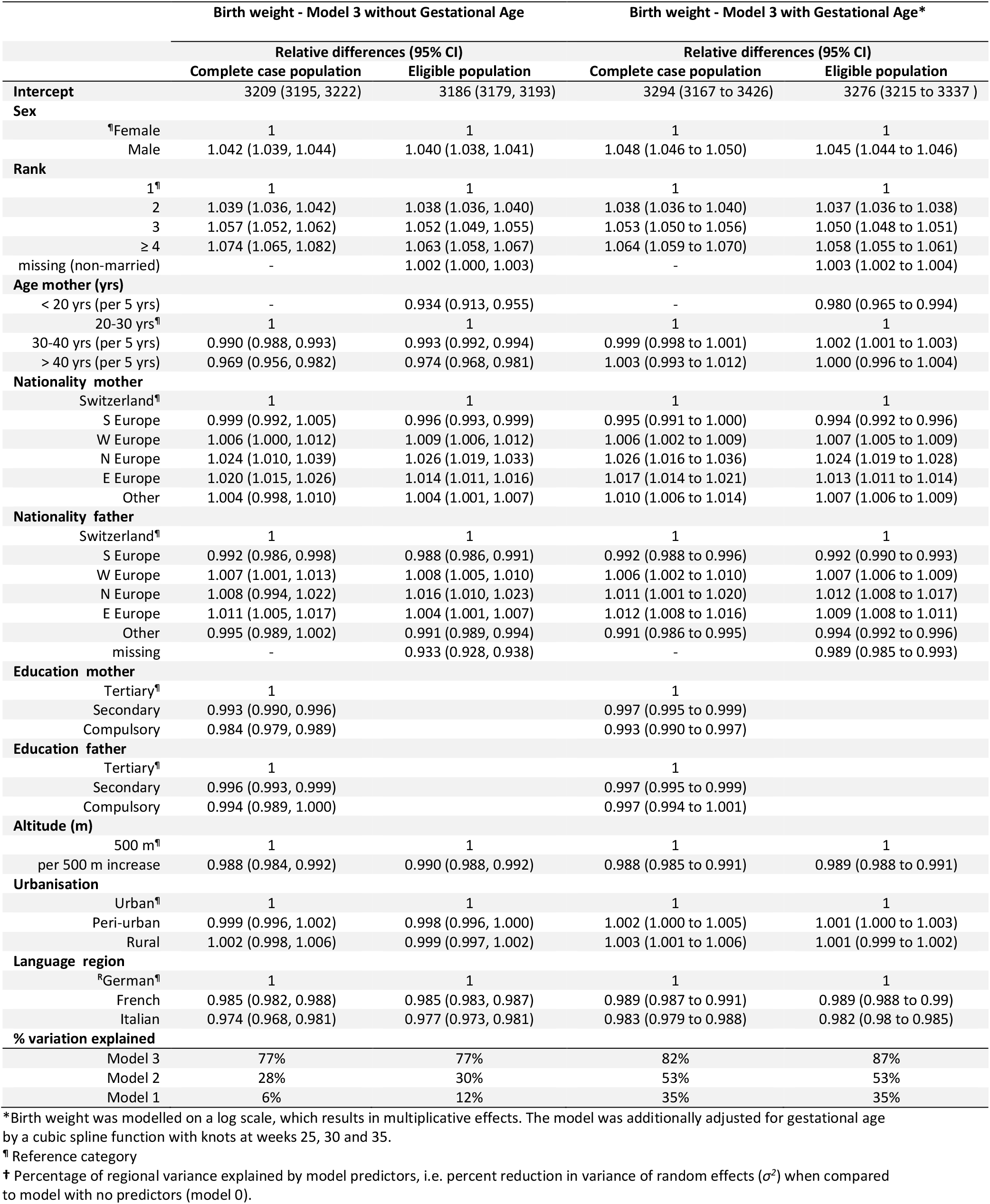
Comparison of results from model (model 3) for birth weight, adjusted and not adjusted for gestational age.

**Supplementary Figure S1.**
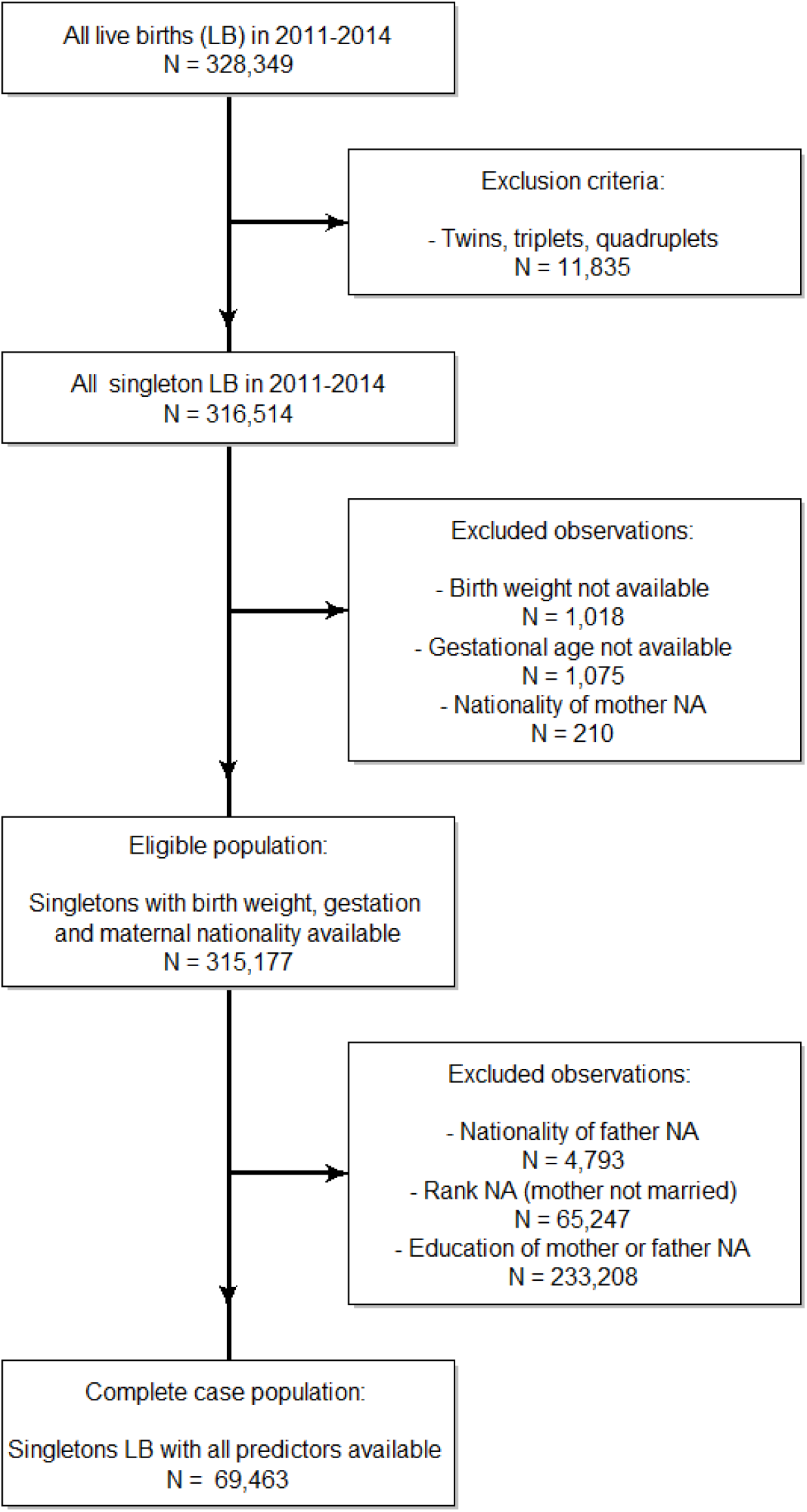
Selection of eligible and complete case study populations among live births in Switzerland 2011 to 2014.

**Supplementary Figure S2.**
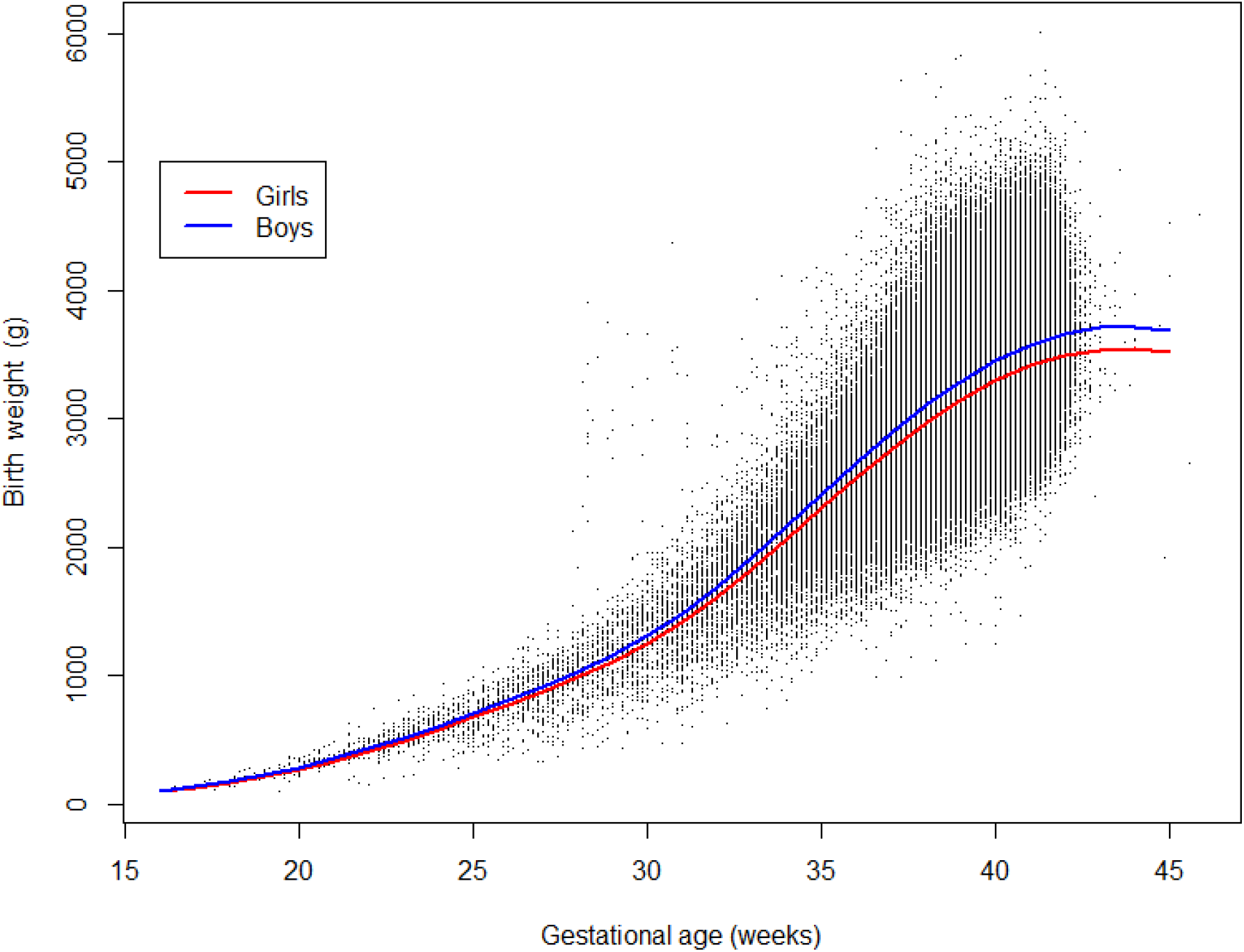
Relationship between birth weight and gestational age at birth modeled by a cubic spline function. Separate fitted curves are shown for newborn girls and boys, with all other predictors corresponding to the reference categories shown in Table 2.

**Supplementary Figure S3.**
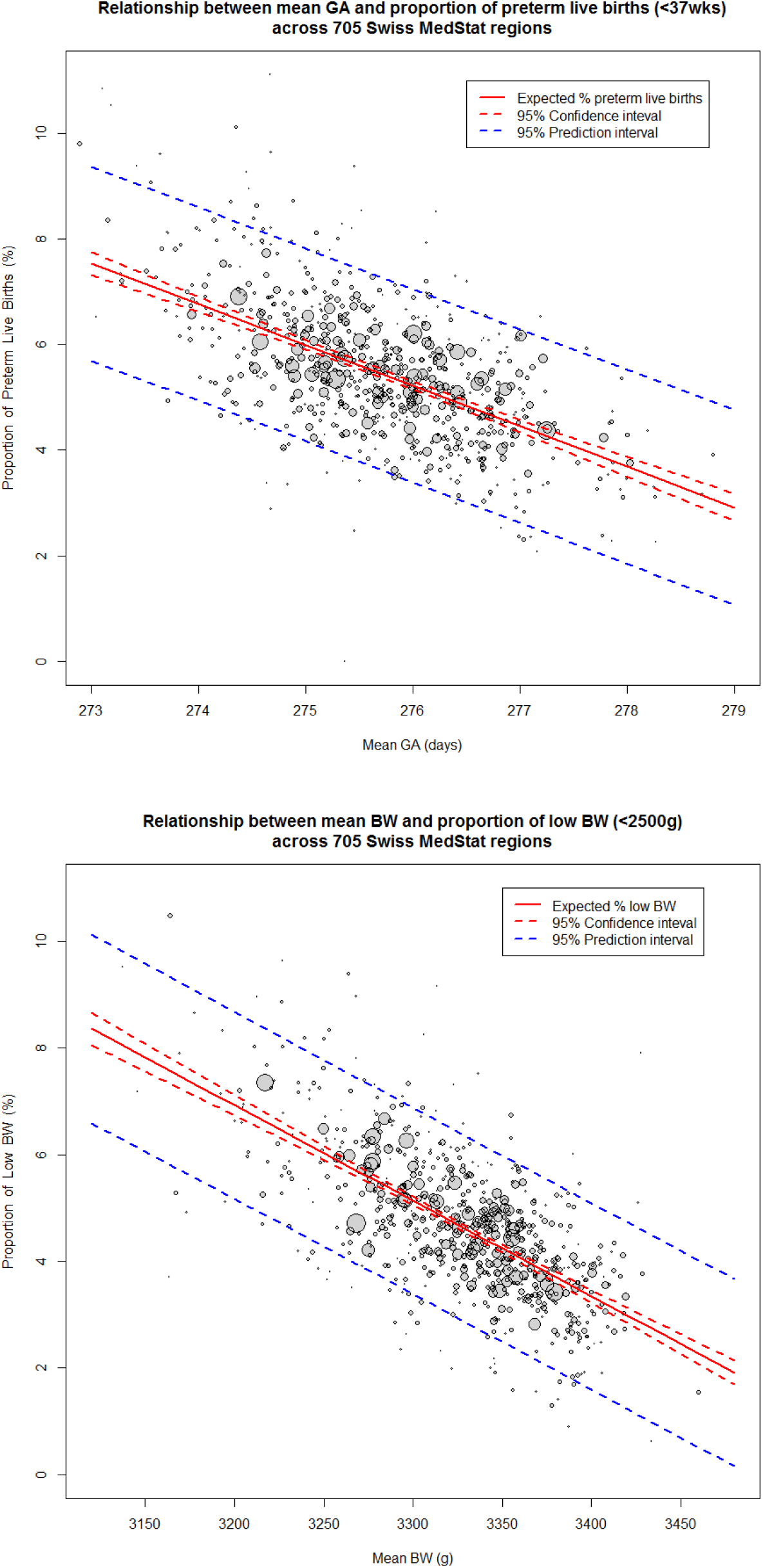
Relationship between mean gestational age and proportion of preterm live births (<37 weeks) among eligible live births across 705 regions (upper panel) and between mean birth weight and proportion of low birth weight births (<2500g) (lower panel). Results from linear regression weighted by the number of live births in each region. Prediction interval displayed for an average-size region (n=447). GA = gestational age; BW= birth weight

## FOOTNOTES

### Contributors

ME and CEK conceived the study and obtained funding. VS, ME, MZ and CEK developed the analysis plan. VS did all statistical analyses and wrote the first draft of the paper, which was revised by ME taking into account the critical comments from all authors. ME supervised the study. All authors approved the final version of the report.

### Patient consent

Not required.

### Ethics approval

The SNC has been approved by the Ethics Committee of the Canton of Bern.

### Provenance and peer review

Not commissioned; externally peer reviewed.

### Data sharing statement

Data are available within the framework of a data sharing agreement.

